# Threshold Electric Skin Sensitivity Fluctuations in Pregnancy, Labor and Puerperium

**DOI:** 10.1101/565960

**Authors:** Charles Boris Cernoch Rossmann, Cosima von Taaffe

**Affiliations:** Department of Obstetrics and Gynecology, Western Memorial Regional Hospital in Corner Brook, Newfoundland, Canada, A2H 6J7

## Introduction

The Threshold Electric Skin Sensitivity (TESS) was intensively investigated by many researchers during the past 100 years and a good survey of the previous literature is discussed in the work of Notermans, 1966 [8]. He used square wave constant current stimulus and tried to determine how many variables could have an influence on the skin sensitivity to electric stimulation. He found nearly constant pain threshold when measured in the same individual over a course of time. Schumacher et al, 1940 [12] measuring threshold sensitivity to heat stimulation in an earlier study came to the same conclusion. On the contrary, Lanier, 1943 [6] found that the threshold sensitivity to electric skin stimulation could vary considerably. Most authors measured the skin sensitivity in many different skin areas, but not repeatedly in short intervals over the same area of the same subject.

Uher et al, 1963 [13] found decreased electric skin sensitivity above the pubic symphysis in early labor and after the administration of oxytocin. He found increased sensitivity after administration of strychnine. In our previous study Cernoch et al, 1969 [2], was used the same stimulator with constant voltage square waves on pregnant and puerperal women and we found long term changes of skin sensitivity. Later, we also found interesting short-term changes of sensitivity when we used a stimulator with constant current square waves on puerperal women after administration of oxytocin and neostigmine. Cernoch et al, 1970 [1]. In pregnant women we found significant skin sensitivity changes after intravenous administration of oxytocin already before the beginning of the uterine contraction. We tried to find, under the same conditions, changes in other biophysical parameters of the skin; however, measurement of the skin temperature and electric conductivity did not reveal any similar changes. Cupr et al. 1972 [3] was using the same constant current square wave stimulator constructed by Valosek et al, 1969 [14]. They found different skin sensitivity during and between the uterine contractions in laboring women with a history of dysmenorrhea and no change in women with no dysmenorrhea.

As we found in the previous investigation Cernoch et al, 1969, 1970 [2,1], there are long term and short-term changes of the Threshold Electric Skin Sensitivity TESS in specific areas of the abdomen during pregnancy, labor, and puerperium. During the labor, the average TESS above the pubic symphysis and in the lateral areas of the lower abdomen is increased but around the navel is decreased. After administration of oxytocin to puerperal women, the usually steady TESS in the lateral abdomen suddenly starts fluctuating up and down. After administration of neostigmine, there is no change of steady TESS in lateral areas; but there is a prominent increase of TESS fluctuations around the navel and above the pubis. These TESS fluctuations may be related to the functional state of the internal organs like uterus and bowel. The projection referred pain skin area for the uterus is on both sides of the lower abdomen, fusing together above the pubic symphysis as a referred pain skin area for the uterine cervix Rubin, 1947 [11]. The referred pain skin area for the bowel is in the midline of the abdomen. Jones, 1938 [5].

## Methods

The Threshold Electric Skin Sensitivity (TESS) was measured repeatedly in short intervals on abdomen and right forearm of 70 pregnant and not pregnant women. The subjects were informed of the measurement procedure and they gave informed consent to participate in the study. The mean age was 23 years (range 15 to 43 years). They have been randomly selected from available patients that came in the clinic and hospital for treatment. Rossmann et al, 1991 [10].

We also used in this study different stimulator, then that was used in previous research. It was an NS-3 Peripheral nerve stimulator from Professional Instruments Co. (Houston, Texas). It produces a rectangular single pulse of 0.2 millisecond duration in one second intervals. The stimulating current can be regulated from 0 to 20 milliamperes and the output voltage can be read from the scale. As a stimulating electrode we used disposable Red Dot 3M Monitoring electrodes with solid gel and Micropore R tape. They were placed on the abdomen and right arm (Figure 1). As a grounding electrode, we used a disposable NDM Día Temp II self-adhering pad placed on the right thigh.

**Figure 1.**
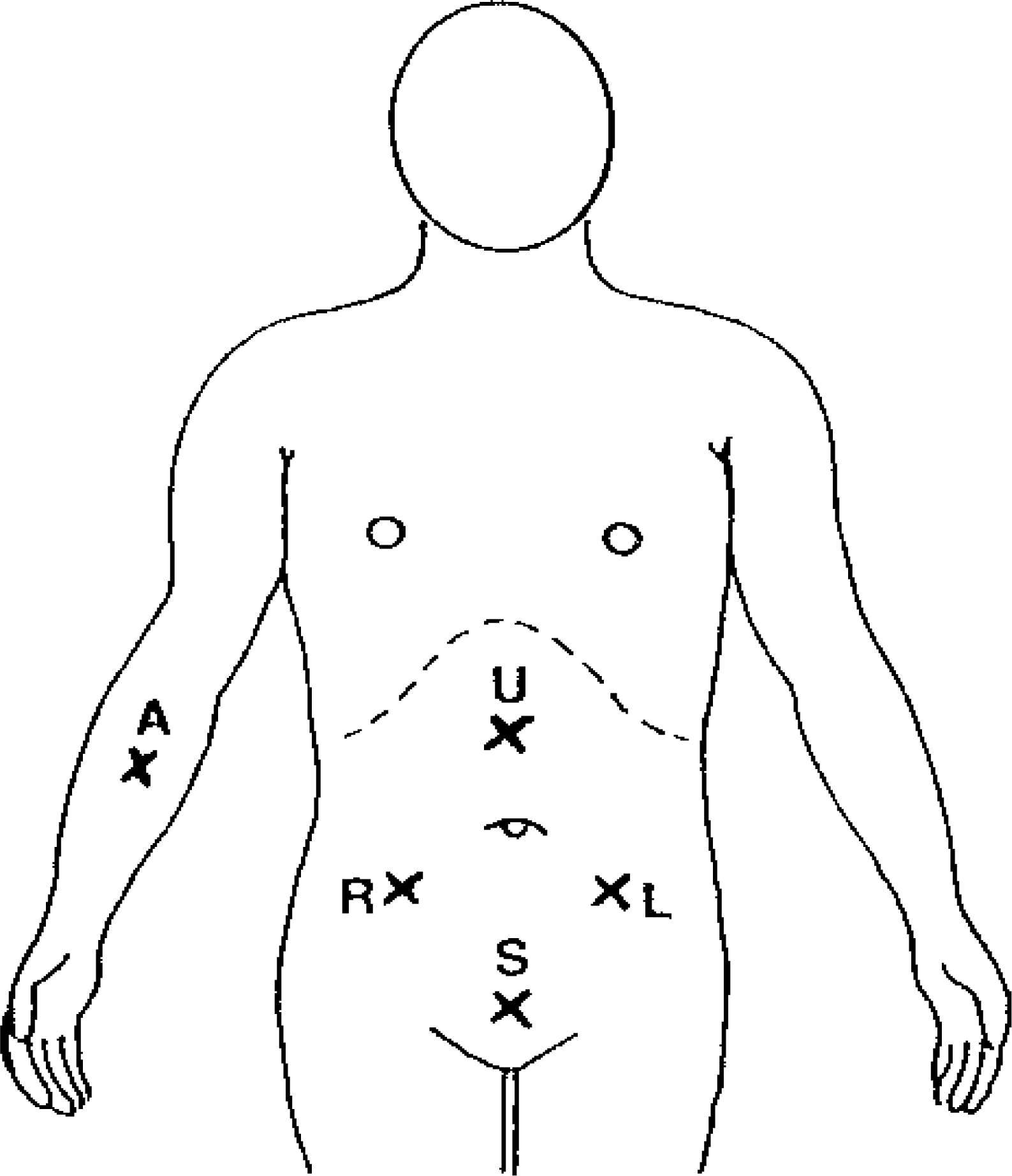
Location of stimulating electrodes on abdomen and forearm: R = Right lower abdomen, L = Left lower abdomen, U = Midline above umbilicus, S = Midline above symphysis, A = Right forearm.

Each subject would lie relaxed on the bed or on the examination table. Before starting the measurement, the patient was instructed to immediately say “yes” when she started to feel the first mild sensation below the electrode. After that, a few orientation stimulations were done to make the patient familiar with the procedure and to exclude the effect of learning on the results. The sequence of measurements was right side of lower abdomen, left side of lower abdomen, middle above umbilicus, middle above pubic symphysis, and on the right forearm. We have done in 5 to 10 second intervals 30,300 single TESS measurements during 1,180 measurement episodes lasting 3 to 5 minutes or longer on each electrode in 280 measurement settings lasting 15 to 30 minutes on 70 patients. During the labor, the patient’s uterine contractions were simultaneously recorded by a Hewlett-Packard cardiotocograph fetal monitor. Uterine contractions and skin TESS fluctuations have been later transcribed in the same graphic record for easy visual correlation. The length of TESS measurements during labor has been limited by the severity of the distress from pain and the willingness of the patient to cooperate. Patients did not receive any analgesia before the measurement.

The TESS measured score data were dictated in to a tape recorder, transcribed in tables as a function of the time and plotted on the graph. To express the TESS fluctuations phenomenon in score data, the length of the curve line was measured in millimeters and divided by the time of measurement in minutes (mm/min.). The higher score represents fluctuations of bigger amplitude or frequency. Later we started using a simpler method for expressing the TESS fluctuations in score data. We calculated the Standard deviation for values of TESS sensitivity collected in each measurement episode. The value of the Standard deviation used as a score data, closely correlate with the score data received by measuring the length of the plotted sensitivity curve per minute. Only score data obtained this way were used for further statistical analysis and presentation in this report. The measured Threshold Electric Skin Sensitivity data were plotted as a function of a time in graph. Figure 2 shows examples of typical Electrosensitogram (ESG) in non-pregnant, pregnant, and post partum women.

**Figure 2.**
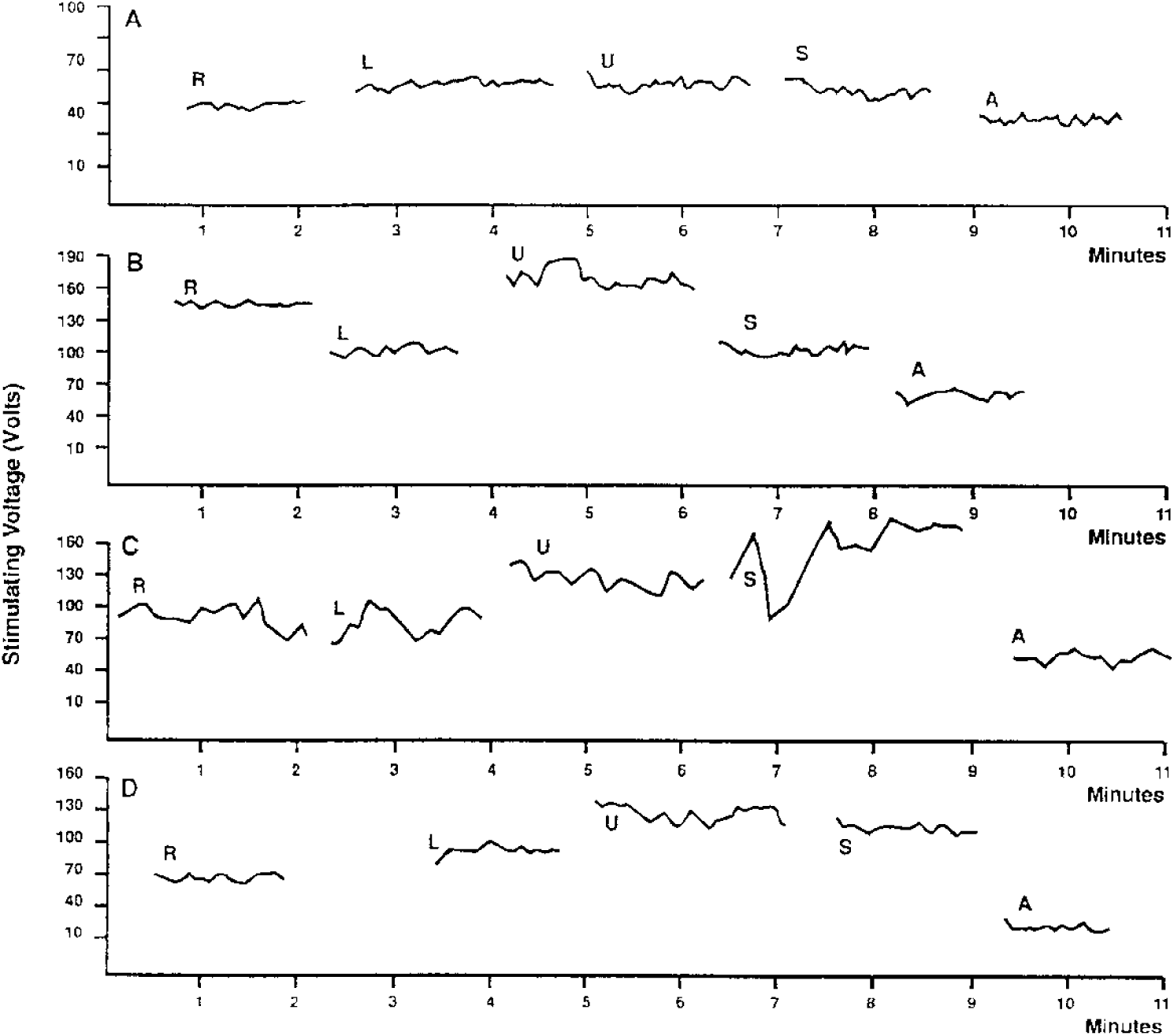
Examples of Electrosensitogram (ESG) graphs: (A) Non-pregnant women), (B) At 29th weeks of pregnancy, (C) At 39th week of pregnancy, (D) Post partum. Location of electrodes: R = Right lower abdomen, L = Left lower abdomen, U = Midline above umbilicus, S = Midline above symphysis, A = Right forearm.

In the presented study, one level of the independent variable under our control was a group of non-pregnant women. The next levels of the independent variable were pregnant or post-partum patients divided in groups according to gestational age, irregular or regular contractions during labor, and days post partum. For all the statistical analysis presented in this report, only two levels of the independent variable were used in One-way, between-subjects analysis of variance (ANOVA). The dependent variable is the calculated score data of TESS fluctuations. The alpha level was selected at 0.05. Each time we found a statistically significant difference between the two levels of independent variable, the Strength-of-association measure test was done using eta-squared computational formula.

## Results

Figure 3 illustrates fluctuations of Threshold Electric Skin Sensitivity TESS in five different areas of the skin of the abdomen and right forearm. The height of the bars indicates the mean intensity of TESS fluctuations in groups of women that are not pregnant, or in consecutive stages of pregnancy and post partum. Above the navel, the sensitivity fluctuates very wildly and inconsistently for most of the time. The only statistically significant change (p < 0.05) is between non-pregnant and pregnant women with irregular contractions. There is a decrease of skin TESS fluctuations when contractions become regular during labor and most of the time, post-partum. Above the pubic symphysis, the TESS fluctuations gradually increase during the pregnancy; but similarly, like above the navel, there is a great variability. Only increases of TESS fluctuations from 36 weeks of pregnancy to the beginning of irregular contractions are statistically very significant (p < 0.01). During regular contractions and post-partum, (like that above the navel) there is a marked decrease of TESS fluctuations.

**Figure 3.**
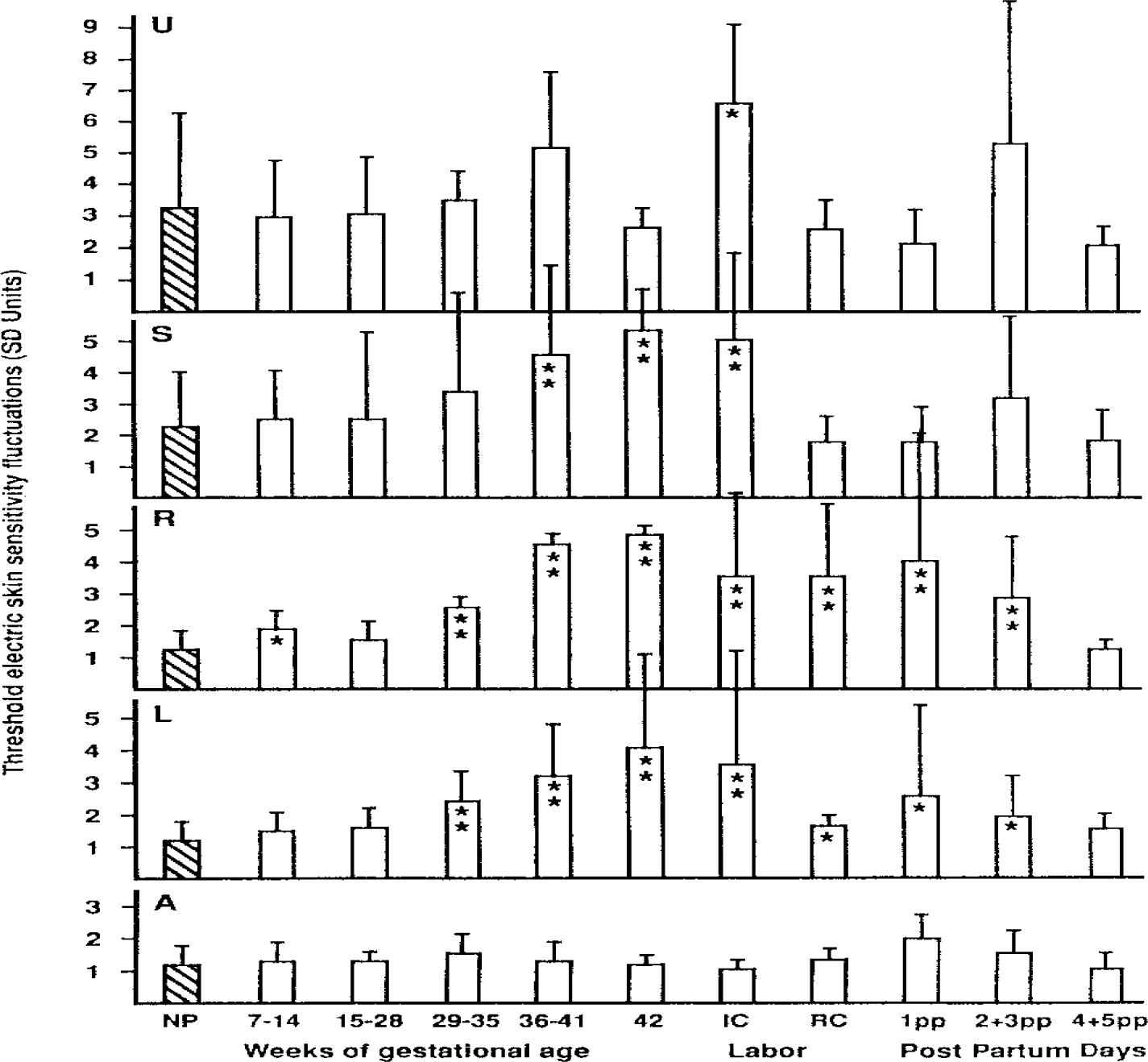
Changes of Threshold electric skin sensitivity (TESS) fluctuations during pregnancy, labor and post partum. PANELS: U = Midline above umbilicus, S = Midline above symphysis, R = Right lower abdomen, L = Left lower abdomen, A = Right forearm. BARS: NP = Not pregnant, 7-14, 15-28, 29-35, 36-41, 42 = Weeks of gestational age, IC = Irregular contractions, RC = Regular contractions, 1pp, 2+3pp, 4+5pp = Days post partum. Values statistically significant are indicated by single (p < 0.05) or double (p < 0.01) asterisk.

Both lateral skin areas of the abdomen behave in a totally different manner to that of the medial areas. There is consistently, only minimal fluctuation of TESS in non-pregnant women and in early pregnancy. But statistically, there is a very significant increase (p < 0.01) of TESS fluctuations from the 29th week of pregnancy to the beginning of irregular contractions. In most of the women, post-partum TESS fluctuations decrease in lateral areas; however, few of them had very big TESS fluctuations and that caused a large variability in the measured data. The control skin area on the right forearm did show consistently minimal fluctuations of TESS and no statistically significant changes between all groups of patients were found. It is interesting that from the fourth day, post-partum, the TESS fluctuations in all areas return to the values of non-pregnant women. In contrast, there was a large fluctuation and variability of TESS on the third day after labor, especially above the navel.

In 10 women, the contractions were recorded simultaneously during the skin TESS measurement. As can be seen on Figure 4A, the tocographic record has been transposed on the skin sensitivity graph so that visual correlation can be done between both curves. We found that women with irregular contractions have more TESS fluctuations than later, during the regular uterine contractions. The means for the levels of the independent variables were reported in Table 1. Analysis of the variance performed on this data indicated that there were statistically significant differences (p < 0.05) among the means for the factors presented in Table 2. The null hypothesis must therefore be rejected, and the conclusion reached that the fluctuations of the Threshold Electric Skin Sensitivity (TESS) changes in certain skin areas of the abdomen during pregnancy, labor and post partum. The Strength-of-association measure (eta-square) for the statistically significant changes with p < 0.05 was between 11% and 22%. The eta-square for statistically very significant changes with p < 0.01 was found between 26% and 62%. It indicates that in these situations there is a very strong relationship between the independent and dependent variables and provides justification for making strong inferences about the validity of the presented results.

**Figure 4.**
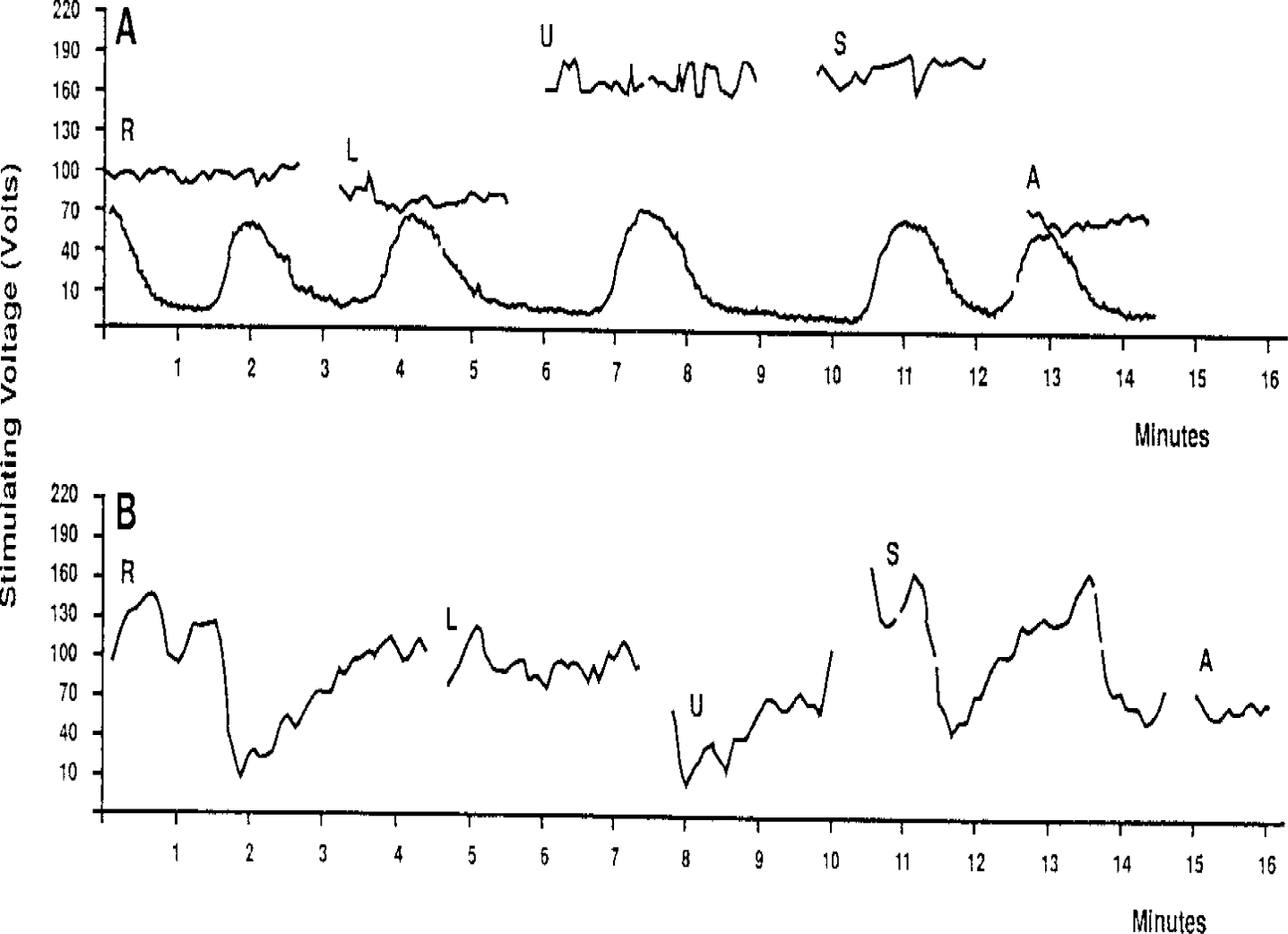
(A) Tocogram recorded simultaneously with Electrosensitogram (ESG) during labor. (B) Electrosensitogram (ESG) of patient with mild preeclampsia at 38 weeks of gestational age.

**Table 1.** Changes of Threshold electric skin sensitivity (TESS) fluctuations during pregnancy, labor and post partum.

| Measured<br>in area | Para-<br>meter | Not<br>preg-<br>nant | Preg-<br>nant<br>week<br>7-14 | Preg-<br>nant<br>week<br>15-18 | Preg-<br>nant<br>week<br>29-35 | Preg-<br>nant<br>week<br>36-41 | Preg-<br>nant<br>week<br>42 | Contrac-<br>tions<br>not<br>regular | Contrac-<br>tions<br>regular | Post<br>partum<br>day<br>1 | Post<br>partum<br>day<br>2+3 | Post<br>partum<br>day<br>4+5 |
| --- | --- | --- | --- | --- | --- | --- | --- | --- | --- | --- | --- | --- |
| Middle<br>above<br>umbilicus | Mean | 3.2 | 3.0 | 3.1 | 3.4 | 5.1 | 2.7 | 6.6 | 2.5 | 2.1 | 5.2 | 2.0 |
|  | SD | 3.1 | 1.8 | 1.8 | 0.9 | 2.5 | 0.6 | 2.6 | 0.9 | 1.1 | 5.6 | 0.6 |
|  | N | 19 | 25 | 5 | 6 | 11 | 4 | 7 | 4 | 4 | 9 | 5 |
| Middle<br>above<br>symphysis | Mean | 2.2 | 2.6 | 2.7 | 3.4 | 4.8 | 5.3 | 5.1 | 1.8 | 1.8 | 3.2 | 2.0 |
|  | SD | 1.8 | 1.5 | 2.7 | 3.2 | 2.7 | 1.2 | 2.6 | 0.7 | 0.8 | 2.7 | 0.8 |
|  | N | 19 | 25 | 5 | 6 | 11 | 4 | 7 | 4 | 4 | 9 | 3 |
| Right<br>lower<br>abdomen | Mean | 1.2 | 1.9 | 1.4 | 2.7 | 4.5 | 4.8 | 3.4 | 3.5 | 4.1 | 2.9 | 1.2 |
|  | SD | 0.6 | 1.3 | 0.3 | 2.1 | 2.8 | 2.6 | 2.4 | 3.3 | 4.1 | 2.0 | 0.4 |
|  | N | 19 | 25 | 5 | 6 | 11 | 4 | 7 | 4 | 4 | 9 | 3 |
| Left<br>lower<br>abdomen | Mean | 1.2 | 1.5 | 1.5 | 2.3 | 3.2 | 4.2 | 3.6 | 1.7 | 2.6 | 1.0 | 1.6 |
|  | SD | 0.5 | 0.6 | 0.7 | 0.8 | 1.7 | 2.9 | 3.5 | 3.6 | 3.0 | 1.2 | 0.5 |
|  | N | 19 | 25 | 5 | 6 | 11 | 4 | 7 | 4 | 4 | 9 | 5 |
| Right<br>forearm | Mean | 1.3 | 1.4 | 1.3 | 1.7 | 1.5 | 1.2 | 1.1 | 1.3 | 2.1 | 1.6 | 2.1 |
|  | SD | 0.6 | 0.5 | 0.1 | 0.6 | 0.5 | 0.3 | 0.3 | 0.4 | 0.8 | 0.8 | 0.5 |
|  | N | 20 | 23 | 3 | 5 | 10 | 3 | 6 | 4 | 3 | 6 | 4 |

Mean = Mean threshold electric skin sensitivity fluctuations.
SD = Standard deviation from mean
N = Number of measured subjects

**Table 2.** One-Way Between-Subjects ANOVA of Threshold Electric Skin Sensitivity (TESS) Fluctuations.

| Source | df | SS | MS | F | Eta square | p | Significance |
| --- | --- | --- | --- | --- | --- | --- | --- |
| Factor R:7-14 GA | 1 | 5.3 | 5.3 | 5.01 | 11% | < 0.05 | * |
| Error | 42 | 44.7 | 1.1 |  |  |  |  |
| Factor R:29-35 GA | 1 | 10.7 | 10.7 | 8.51 | 27% | < 0.01 | ** |
| Error | 23 | 29.1 | 1.3 |  |  |  |  |
| Factor L:29-35 GA | 1 | 5.5 | 5.5 | 17.25 | 43% | < 0.01 | ** |
| Error | 23 | 7.3 | 0.3 |  |  |  |  |
| Factor S:36-41 GA | 1 | 46.7 | 46.7 | 9.88 | 26% | < 0.01 | ** |
| Error | 28 | 132.5 | 4.7 |  |  |  |  |
| Factor R:36-41 GA | 1 | 76.1 | 76.1 | 24.53 | 47% | < 0.01 | ** |
| Error | 28 | 86.9 | 3.1 |  |  |  |  |
| Factor L:36-41 GA | 1 | 28.4 | 28.4 | 25.14 | 47% | < 0.01 | ** |
| Error | 28 | 31.4 | 1.1 |  |  |  |  |
| Factor S:42 GA | 1 | 32.6 | 32.6 | 10.98 | 34% | < 0.01 | ** |
| Error | 21 | 62.4 | 3.0 |  |  |  |  |
| Factor R:42 GA | 1 | 42.8 | 42.8 | 34.42 | 62% | < 0.01 | ** |
| Error | 21 | 26.1 | 1.2 |  |  |  |  |
| Factor L:42 GA | 1 | 29.1 | 29.1 | 20.75 | 50% | < 0.01 | ** |
| Error | 21 | 29.5 | 1.4 |  |  |  |  |
| Factor U: IC | 1 | 60.3 | 60.3 | 6.86 | 22% | < 0.05 | * |
| Error | 24 | 210.9 | 8.8 |  |  |  |  |
| Factor S: IC | 1 | 44.4 | 44.4 | 10.69 | 31% | < 0.01 | ** |
| Error | 24 | 99.6 | 4.2 |  |  |  |  |
| Factor R: IC | 1 | 24.9 | 24.9 | 14.78 | 38% | < 0.01 | ** |
| Error | 24 | 40.4 | 1.7 |  |  |  |  |
| Factor L: IC | 1 | 30.8 | 30.8 | 9.53 | 28% | < 0.01 | ** |
| Error | 24 | 77.6 | 3.2 |  |  |  |  |
| Factor R:RC | 1 | 18.0 | 18.0 | 16.93 | 45% | < 0.01 | ** |
| Error | 21 | 22.3 | 1.1 |  |  |  |  |
| Factor L:RC | 1 | 0.9 | 0.9 | 4.70 | 18% | < 0.05 | * |
| Error | 21 | 4.2 | 0.2 |  |  |  |  |
| Factor R:1pp | 1 | 27.6 | 27.6 | 10.23 | 33% | < 0.01 | ** |
| Error | 21 | 56.7 | 2.7 |  |  |  |  |
| Factor L:1pp | 1 | 7.0 | 7.0 | 4.86 | 19% | < 0.05 | * |
| Error | 21 | 30.1 | 0.3 |  |  |  |  |
| Factor R:2+3pp | 1 | 17.8 | 17.8 | 11.71 | 31% | < 0.01 | ** |
| Error | 26 | 39.5 | 1.5 |  |  |  |  |
| Factor L:2+3pp | 1 | 4.3 | 4.3 | 7.67 | 22% | < 0.05 | * |
| Error | 26 | 14.7 | 0.6 |  |  |  |  |

U = Midline above umbilicus.
S = Midline above symphysis.
R = Right lower abdomen.
L = Left lower abdomen.
GA = Gestational age weeks
IC = Irregular contractions in labor.
RC = Regular contractions in labor.
df = Degree of freedom
SS = Sum of squares

Measurements have been also done on several patients with preeclampsia, severe gestationaledema or irritable bowel, and it did show especially large TESS fluctuations in most of the measured areas (Figure 4B). Similar, large TESS fluctuations of sensitivity were found in one 34-week pregnant patient with acute functional ureteric obstruction. After the obstruction was relieved by ureteric stent, the TESS fluctuations gradually decreased over several days (Figure 5).

**Figure 5.**
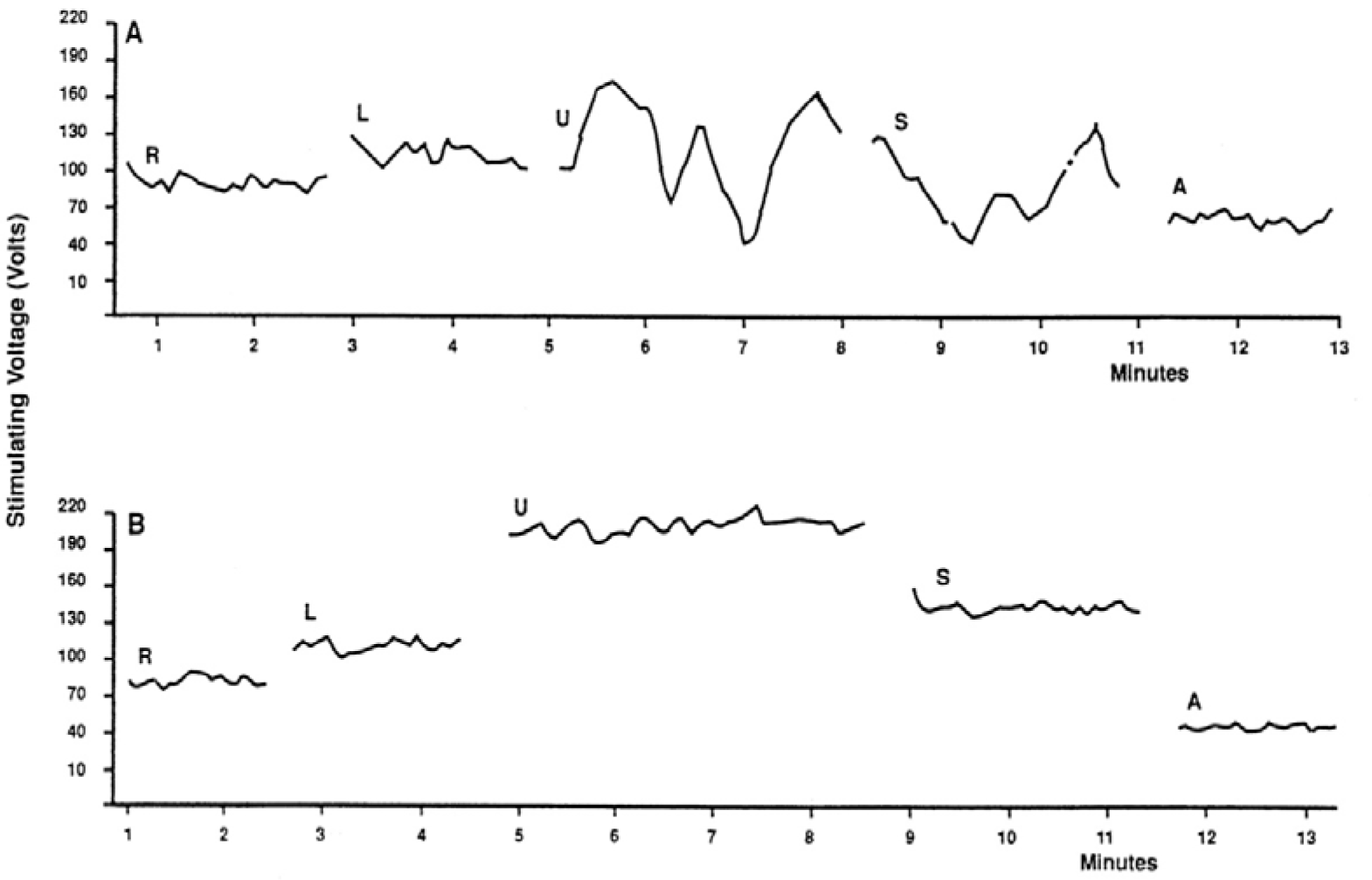
(A) Electrosensitogram (ESG) of 34 weeks pregnant patient with acute functional ureteric obstruction. (B) Same patient four days after the obstruction was relieved by ureteric stent.

## Discussion

Our present work supports the research hypothesis that there are changes of short-term Threshold electric Skin Sensitivity fluctuations during consecutive stages of pregnancy, labor, and post partum. Maximal TESS fluctuations were found in all measured skin areas during late pregnancy and during irregular contractions at the beginning of labor. During regular uterine contractions however, the TESS fluctuations decreased, and we could not confirm our research hypothesis that there would be some temporal relationship between the sensitivity curve and the tocographic record of the uterine contraction. Minimal TESS fluctuations were generally found in most of the skin areas of non-pregnant women and this served us as a control independent variable. Increased TESS fluctuations during late pregnancy and labor are consistent with our earlier research findings, when we administered oxytocin and neostigmine to puerperal women. The underlying neuro-physiological mechanism however is not clear.

As we postulated in previous papers Cernoch et al, 1969, 1970 [2, 1], the skin sensitivity fluctuations in the midline of the abdomen may be somehow related to the functional state (motility) of the bowels. If any such relationship could be proved by further research, it could have practical application in gastroenterology as a non-invasive diagnostic method. Iovino et al [4] found in irritable bowel patients somatic hypoalgesia to electrical stimuli.

The fluctuations of TESS in the lateral abdominal areas, and above the pubic symphysis are more difficult to explain. Oxytocin and labor clearly have a significant effect on it; but we found that the changes were not synchronous with the uterine contractions. On the other hand, we had found in our previous work that TESS fluctuations on the right- and left-side of abdomen change synchronically after giving oxytocin to puerperal women. Cernoch et al, 1970 [1]. The TESS fluctuations in lateral areas of the abdomen may be related to the activity of the autonomous nerve system in the uterus. If it is true, the change of TESS in dermatome would be an expression of the autonomous system activity of the internal organs in the corresponding viscerotome. Because there is an overlapping between the viscerotome of the lower gastrointestinal tract and the internal genital system in women, the interpretation of the Electrosensitogram (ESG) may be difficult. Our statistically very significant results indicate that it is not impossible. The anecdotal findings of interesting TESS fluctuations changes in pregnant patients with ureteric obstruction, preeclampsia and irritable bowel syndrome suggest direction for further studies in this and other clinical situations. Other improvement would be combining TESS measurements with objective method to monitor subject’s response to quantitative stimuli. Le et al, 2005 [7]. When we used in previous study constant current stimulator Valosek et al, 1969 [14], based on Laufberger’s excitation theory (Radil) [9], we observed the same short-term fluctuations of TESS Cernoch et al, 1970 [1]. In our now presented investigation we found, that Threshold Electric Skin Sensitivity (TESS) fluctuates in different areas of the abdomen and these fluctuations also change during pregnancy, labor and puerperium. Clinicians are using the manifestations of referred pain in dermatome to help diagnose pathological and physiological processes of the internal organs in the corresponding viscerotome inside the human body. Similarly, the exact measurement and recording of Threshold Electric Skin Sensitivity in dermatome could have diagnostic value. As a non-invasive method, it could help better understand the function of involved internal organs in the corresponding viscerotome.

## Acknowledgments

The authors thank the patients and healthy volunteers who participated in this study.

## Abbreviations

ESG: Electrosensitogram
TESS: Threshold Electric Skin Sensitivity

